# Elemental allocation to molecular drivers of biogeochemistry in the Southern Ocean

**DOI:** 10.1101/2025.10.03.680396

**Authors:** Loay J. Jabre, Elden Rowland, Charlotte Eich, Mathijs van Manen, Corina P.D. Brussaard, Rob Middag, Erin M. Bertrand

## Abstract

Metabolic processes underpinning ocean biogeochemistry are powered by molecular machines, proteins, that require various elements to function. Yet, the allocation of elements to these proteins, and subsequent implications for biogeochemical processes, remain poorly characterized. Here we integrate elemental measurements with metaproteomics to quantitatively examine elemental use in Southern Ocean microbial proteins and metabolic processes. We demonstrate that iron availability influences elemental allocation, including decreased iron allocation to photosynthesis and compensatory incorporation of non-iron metals into metalloproteins under iron scarcity. Manganese was primarily allocated to photosynthesis in iron-replete conditions, and reallocated to other metabolic roles under low iron. Photosystem I:II protein mass ratios impacted both iron and manganese allocation, and appeared to be driven by iron availability. Approximately half of biogenic copper was found in plastocyanin, likely substituting for iron-containing cytochromes in photosynthesis. Moreover, biogenic nitrogen to phosphorus ratios were decoupled from ribosomal abundance, contrary to prevailing assumptions about ribosomal influence on stoichiometric regulation in the ocean. Instead, our results suggest that community composition and intracellular storage are important regulators of N:P in the Southern Ocean. Together, our findings identify key molecular mechanisms that modulate elemental demand and limitation, and provide a foundation for quantitatively connecting molecular measurements with biogeochemical models.

## Main text

Proteins drive essential metabolic processes within ocean microbes, including photosynthesis, respiration, nitrogen fixation, and nutrient uptake. Collectively, these processes shape marine ecosystems and biogeochemical cycles by influencing the biological carbon pump^1–3^, regulating nutrient cycling^4^, and driving food web dynamics^5,6^. We currently lack a predictive, mechanistic understanding of how ocean microbes allocate various elements to the vital metabolic functions they perform. Resolving this uncertainty is essential for constraining biological interactions with elemental gradients in the ocean, and for quantifying microbial influences on biogeochemistry, now and into the future^7,8^.

Proteins represent a large fraction of cellular nitrogen quotas due to their amino acid composition^9^, and because 30-50% of proteins require metal cofactors to function (i.e. metalloproteins)^10,11^, they also constitute the dominant intracellular reservoir of trace metals^12–15^. Similarly, the high abundance of ribosomal protein-RNA complexes, which are phosphorus rich, makes them a key component of intracellular phosphorus pools^16^. Proteomic analyses of marine microbes can therefore provide insights into elemental allocation to biogeochemically-relevant metabolic processes. Despite this, no studies have directly quantified elemental allocation to biogeochemical functions in marine systems primarily due to logistical and methodological challenges, and only a few have investigated element-protein couplings in laboratory cultures of marine microorganisms^17,18^.

Here we combined metaproteomic profiling with elemental measurements to investigate the allocation of critical elements to specific protein classes in natural Southern Ocean (SO) microbial assemblages. This revealed previously unrecognized patterns in elemental use across key metabolic processes, providing insights into the molecular drivers of stoichiometric variability in one of Earth’s key ocean systems.

### Shipboard incubations of two distinct microbial assemblages in the Weddell Sea

The Weddell Sea in the SO plays a vital role in exporting heat and carbon to the deep ocean, and modulates nutrient delivery to lower latitudes^19–21^. Primary productivity in this region is often constrained by low light, temperature, and iron availability^22–24^, all of which exhibit seasonal variability and are predicted to change under future climate scenarios^25^. Iron is an essential element for most life as it is involved in redox catalysis and electron transport of many metabolic processes including photosynthesis, respiration and nitrogen fixation^26–28^. Temperature directly influences enzyme kinetics and membrane fluidity^29^, ultimately regulating growth and physiology^30^, and may interactively, with iron, influence growth of SO phytoplankton^31–33^.

Here we conducted two shipboard iron-temperature manipulation experiments at two locations in the Weddell Sea (Offshore and Nearshore, Fig. 1A, Fig. S1) during the austral summer of 2018-2019 (Table 1)^33^. Water was collected using an ultraclean CTD sampling system^33–35^, and incubated for eight days under the different treatments (Table 1) before harvesting for metaproteomics and elemental measurements (see Methods). Both locations were characterized by iron limitation, but more so the Offshore location, as illustrated by in-situ molar ratios of dissolved (indicating supply) to particulate iron (Fig. 1B). This is consistent with other signatures of iron limitation in these experiments, including suppressed growth and nutrient drawdown, which were stimulated upon iron enrichment^33^. The Offshore location also had low in-situ manganese supply (Fig. 1B, Fig. S2), indicating possible manganese impacts on phytoplankton growth and community composition^36,37^. All other nutrients were expected to be replete at both locations (Fig. 1B).

**Figure 1.**
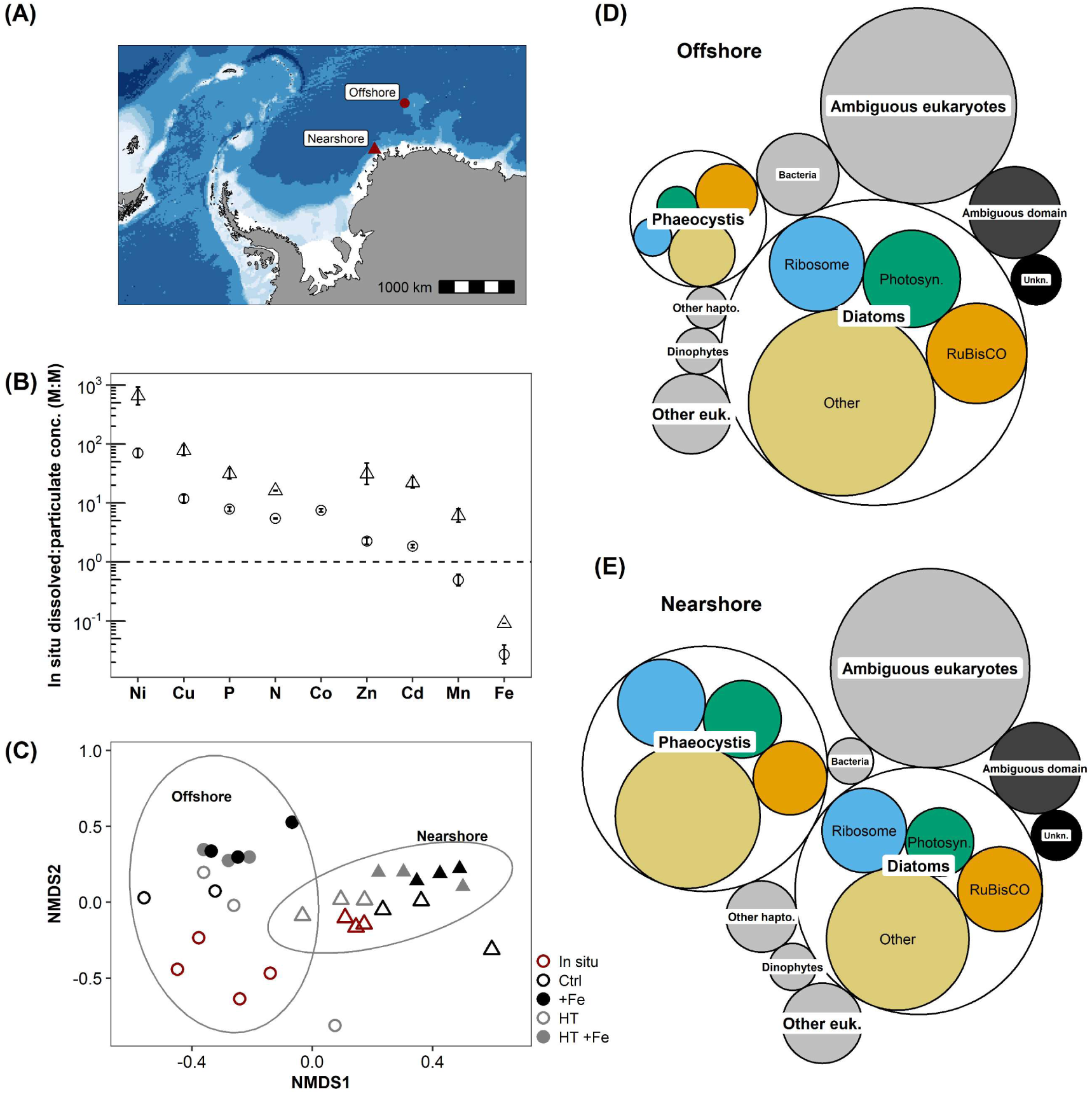
Distinct elemental profiles and microbial assemblages at the Offshore and Nearshore locations. **A)** Locations of the Offshore (circle - 65 °S, 0.038 °E) and Nearshore (triangle - 70.15 °S, 11.02 °W) study sites in the Weddell Sea, Southern Ocean. **B)** In-situ ratios of dissolved to particulate (biogenic) elemental concentrations (M:M). Horizontal dashed line marks a 1:1 ratio, values above this line indicate higher supply than demand (replete) and values below the line indicate lower supply than demand (limitation). Points represent means of triplicate measurements, error bars represent ± 1SD and fall within the bounds of points when not visible. **C)** Non-metric multi-dimensional scaling (NMDS) ordination based on peptide abundance profiles in-situ (red) and after iron-temperature incubations. Open and closed symbols denote no iron added and iron added treatments, respectively. Black and grey colors denote no warming and warming treatments, respectively. NMDS stress value = 0.12. Each point represents one biological replicate. In B) and C), circles and triangles indicate Offshore and Nearshore samples, respectively. **D, E)** Circle plots showing in-situ abundance of major taxonomic and functional protein groups Offshore (D) and Nearshore (E). Blue = ribosomes, green = light reaction photosynthetic proteins, orange = RuBisCO. Circle size reflects taxon-specific peptide abundance normalized to total matched peptide abundance at each site. “Unkn” represents peptides with no known function or taxonomic affiliation, “Ambiguous eukaryotes” represents peptides mapped to multiple eukaryotic taxa (e.g. peptide sequence found in both diatoms and *Phaeocystis*), “Ambiguous domain” represents peptides mapping to multiple domains (e.g. peptide sequence found in both eukaryotes and bacteria).

**Table 1.** Factorial design of the shipboard iron-temperature incubations. The four different treatments were chosen to mimic current and projected iron-temperature conditions in the Weddell Sea^44–47^: *i*) a control of in-situ temperature and no iron addition (*Ctrl*), *ii*) iron (^57^Fe) addition at in-situ temperature (*+Fe*), *iii*) warming—high temperature—with no iron addition (*HT*), *iv*) concurrent warming and iron (^57^Fe) addition (*HT +Fe*). The shipboard incubators could not consistently maintain Nearshore in-situ temperature of −1.4 °C, so we increased the temperature by 0.4 °C for all treatments in this experiment (see Methods and Eich and van Manen et al. 2024^33^ for details).

|  | Treatment | Temperature | Iron |
| --- | --- | --- | --- |
| **Offshore** | Ctrl | -0.3 °C (in-situ) | 0.05 nM <sup>56</sup>Fe (in-situ) |
|  | +Fe | -0.3 °C (in-situ) | +2 nM <sup>57</sup>Fe |
|  | HT | 1.7 °C (in-situ +2 °C) | 0.05 nM <sup>56</sup>Fe (in-situ) |
|  | HT +Fe | 1.7 °C (in-situ +2 °C) | +2 nM <sup>57</sup>Fe |
| **Nearshore** | Ctrl | -1.0 °C (in-situ +0.4 °C) | 0.03 nM <sup>56</sup>Fe (in-situ) |
|  | +Fe | -1.0 °C (in-situ +0.4 °C) | +2 nM <sup>57</sup>Fe |
|  | HT | 1.0 °C (in-situ +2.4 °C) | 0.03 nM <sup>56</sup>Fe(in-situ) |
|  | HT +Fe | 1.0 °C (in-situ +2.4 °C) | +2 nM <sup>57</sup>Fe |

We used metaproteomics to assess microbial community composition and functional characteristics. We first mapped individual peptides to corresponding taxonomic and functional annotations, then pooled peptides by their annotations into taxonomic and functional groups (see Methods). A non-metric multidimensional scaling ordination at the peptide level identified two distinct peptide clusters, corresponding to microbial assemblages Offshore and Nearshore (Fig. 1C). This analysis also showed that iron supplementation induced a more pronounced change in proteome profiles compared to warming (Fig. 1C). We then calculated the abundance of peptides allocated to specific taxonomic groups as a percentage of total peptide abundance (i.e., summed abundance of peptides uniquely matched to taxon*_x_* / total peptide abundance). *Phaeocystis* spp. was more abundant Nearshore (22-40% of total protein) compared to Offshore (8-12% of total protein), whereas diatoms were more abundant Offshore (30-52% of total protein) compared to Nearshore (21-33% of total protein) (Fig. 1D,E). These patterns were consistent with pigment measurements^33^, and align with previously documented partitioning in SO phytoplankton communities, where *Phaeocystis* tends to thrive in colder conditions where trace metals are relatively more replete, while diatoms tend to prevail in relatively warmer and stratified waters^31,38–42^.

Warming alone (+2 °C) had negligible impact on phytoplankton growth and nutrient consumption in our experiments^33^. However, we note that previous studies in other SO regions, which tested more pronounced warming (> +3 °C), have reported significant temperature effects on iron-limited phytoplankton growth, nutrient drawdown, and molecular responses^31,43^. Additionally, we observed that a concurrent iron and temperature increase led to synergistic effects on growth and nutrient consumption, discussed in a recent publication^33^. Here, we primarily focus on connecting elemental and proteomic responses to changing iron availability, as impacts of these small temperature differences, at the proteomic level, were comparatively subtle (Fig. 1C).

### Flexible metabolic strategies for mitigating iron limitation

We observed increased incorporation of non-iron metals into the microbial community under low iron conditions^33^, indicating community shifts towards reduced iron requirement and increased use of non-iron metalloproteins (Fig. 2A-F). We examined protein expression patterns to assess whether these elemental changes are reflected at the molecular level.

**Figure 2.**
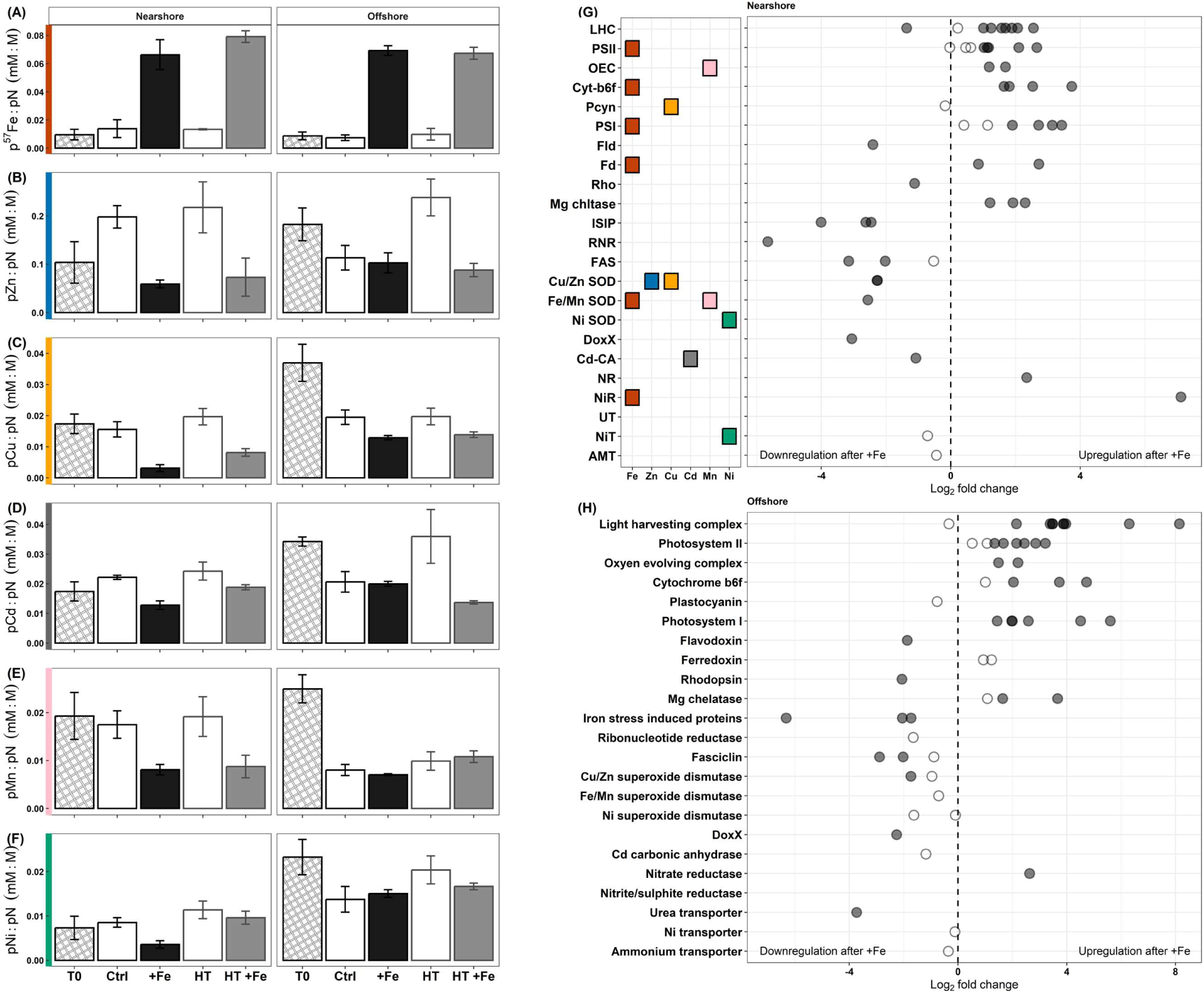
Increased assimilation of non-iron metals into biogenic material under low iron conditions corresponds to elevated abundance of proteins containing these metals. **A-F)** Molar ratios of particulate iron (supplemented ^57^Fe), zinc, copper, cadmium manganese and nickel to particulate nitrogen, respectively. Bar plots show means of triplicate values, error bars represent *±*1 SD and fall within the bounds of bars when not visible. **G, H)** Differential expression of various proteins and protein groups after iron addition at the Nearshore and Offshore locations, respectively. Filled circles represent statistically significant differential expression (adjusted P-value < 0.05). Positive Log_2_ fold change represents increased abundance after iron addition (or decreased abundance under low iron), negative Log_2_ fold change represents decreased abundance after iron addition (or increased abundance under low iron). In G, colored boxes indicate known metal cofactors for each metalloprotein. In H, the Y-axis shows full protein names abbreviated in G. In G and H, each point represents one unique protein or protein group (see Methods). Note that not all proteins were detected and quantified at both study locations.

*Reduced iron requirements* – We measured greater abundance of non-metalated proteins including flavodoxin and rhodopsin under low iron (Fig. 2 G,H), suggesting an increase in metal-free metabolic pathways. While less efficient, flavodoxin can functionally substitute for iron-containing ferredoxin^48^, and rhodopsins may facilitate light-driven energy capture or nutrient uptake, providing benefits in iron-deplete environments^49,50^. We also calculated photosystem I to photosystem II protein mass ratios (PSI:PSII hereafter) directly from the metaproteomics data. These fell within reported ranges measured using alternative methods like photophysiology^51–53^, highlighting our method’s potential for linking protein-level expression with photosynthetic strategies in the field. We observed a marked reduction in PSI:PSII under iron-limited conditions, both in our field experiments and additional laboratory cultures of the Antarctic diatom *Fragilariopsis cylindrus* (Fig. 3A-C). Since PSI requires 3-6 times more iron than PSII^26,54^, this decrease reflects an iron-conserving strategy and shows that in addition to large light-harvesting antennae^53^, SO phytoplankton can minimize iron requirements through tunable photosystem architecture. Lower PSI:PSII may however increase photosynthetic manganese demand due to high manganese requirements in the oxygen-evolving complex (OEC) of PSII, potentially leading to secondary manganese limitation^37^. While our field measurements were made over a relatively small range of manganese conditions, our cultures showed elevated PSI:PSII under low manganese relative to replete conditions, indicating a manganese-conserving response under high iron (Fig. 3B). Further, PSI:PSII and the abundance of manganese-rich PSII proteins appeared to be primarily driven by iron availability in the field (Fig. 3D-F). Future field experiments testing higher ranges of manganese concentrations will allow us to better understand manganese’s influence on phytoplankton photosynthetic processes in the SO.

**Figure 3.**
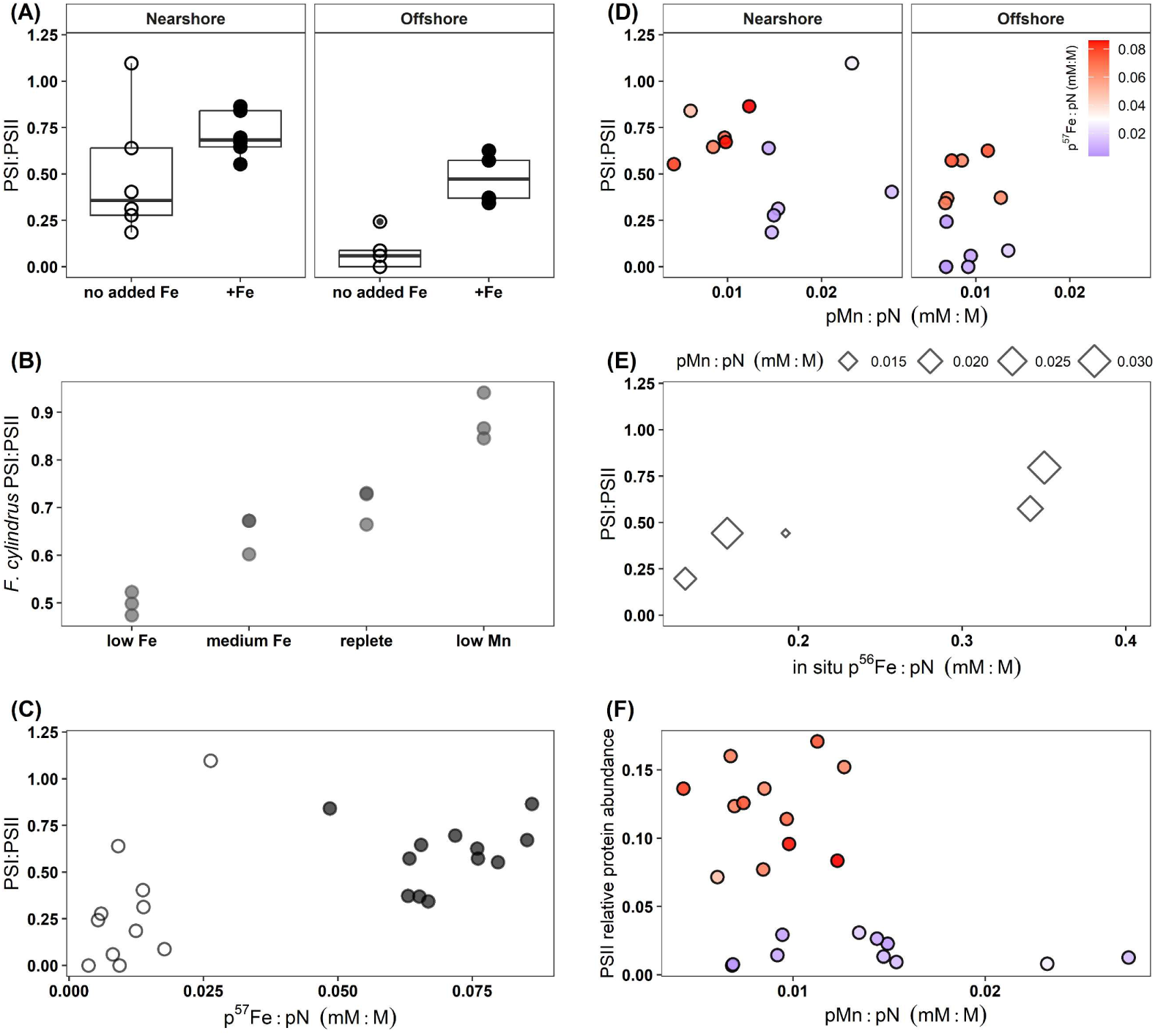
Iron availability influences PSI:PSII mass ratios in natural phytoplankton communities and in cultures. **A)** Photosystem I to photosystem II protein mass ratio (PSI:PSII) calculated as the sum of all PSI proteins divided by the sum of all PSII proteins. Boxplots show median values and upper and lower quartiles. **B)** PSI:PSII measured from *Fragilariopsis cylindrus* laboratory cultures^63^. **C)** PSI:PSII ratios in both Nearshore and Offshore incubation experiments plotted against particulate ^57^Fe (supplemented iron):pN. Filled circles represent iron addition treatments, open circles represent no iron added treatments. **D)** PSI:PSII plotted against particulate manganese (pMn): particulate nitrogen (pN). Point color corresponds to p^57^Fe:pN ratio, where high values (warm colors) correspond to iron addition treatments. **E)** In-situ PSI:PSII plotted against naturally occurring particulate ^56^Fe:pN. **F)** PSII protein abundance, calculated at the sum of all PSII proteins normalized to total protein, plotted against particulate manganese (pMn):particulate nitrogen (pN). Point color corresponds to p^57^Fe:pN ratio as in panel D. In all panels, each point represents one biological replicate.

Upon iron supplementation, we observed increased accumulation of biogenic iron (^57^Fe) (Fig. 2A), accompanied by upregulation of iron metalloproteins including components of the photosynthetic apparatus—PSI, PSII, cytochrome-b_6_f, ferredoxin—and enzymes involved in nitrogen metabolism (Fig. 2G,H). These proteins represent major sinks for intracellular iron in phytoplankton, and their dynamic regulation further underscores the ability of SO phytoplankton to modulate metabolic iron requirements in response to fluctuating iron availability.

*Compensatory use of non-iron metalloproteins –* Under iron-limited conditions, microbial communities increased uptake of non-iron metals, measured as elevated particulate (non-iron) metal-to-nitrogen ratios (pMetal:pN; Fig. 2B-F). Some of this uptake may occur incidentally as iron acquisition pathways, which can also transport other elements, are upregulated under low iron. However, concurrent upregulation of metalloproteins utilizing these metals (Fig. 2G,H) is indicative of active, compensatory metabolic regulation to mitigate iron limitation. For example, particulate copper (pCu:pN) and zinc (pZn:pN) were elevated under low iron (Fig. 2B,C), indicating increased utilization of these metals. Plastocyanin, a copper-binding protein that functionally replaces iron-binding cytochrome-c_6_^55^, was constitutively expressed in our study (Fig. 2G,H), consistent with other observations in the SO^43^. This increase in copper quota under low iron suggests additional uses in other metabolic processes like respiration, iron uptake^56,57^, and increased use in Cu/Zn superoxide dismutase (SOD)^58^, a protein that was generally more abundant under low iron in our study. Similarly, elevated cadmium (pCd:pN) under low iron correlated with increased expression of Cd-carbonic anhydrase (Cd-CA), the only known protein that uses cadmium^59,60^.

Under low iron, biogenic manganese assimilation was elevated at the Nearshore site, whereas the expression of the manganese-rich OEC of PSII was downregulated (Fig. 2E,G). Given that the OEC is a major manganese sink in phytoplankton^14,26^, these findings indicate manganese reallocation from photosynthetic machinery to other metabolic functions under low iron. Notably, many phytoplankton species rely on manganese-binding SOD (Mn-SOD) to manage oxidative stress and other metabolic functions^61^. Here we observed downregulation of Mn-SOD under high iron at the Nearshore location, consistent with previous studies showing increased Mn-SOD use in iron stressed temperate diatoms^62^. Interestingly, this iron-responsive Mn-SOD expression was absent in the Offshore experiment, implicating factors beyond iron availability in influencing its expression: limited manganese availability Offshore (Fig. 1B, Fig S2) may constrain the metalation and use of this enzyme, as shown in a recent study highlighting manganese availability as a key driver of Mn-SOD expression^58^. It is therefore also possible that the Mn-SOD downregulation observed in the Nearshore experiment was driven by manganese being drawn down to limiting concentrations in the high iron treatments, rather than increased iron availability.

### Trace metal allocation to biogeochemically important proteins and metabolic functions

Integrating metaproteomics with elemental measurements can link molecular processes to elemental demand and utilization, thereby providing mechanistic insights into microbial interactions with biogeochemistry. Here we implemented an intensity-based untargeted metaproteomics workflow for quantifying proteins without spike-in standards. Our work builds on previously established methods for protein quantification using label-free shotgun proteomics^64–66^, and extends their application to complex marine microbial communities. By leveraging known protein-metal stoichiometries, we estimated trace metal quotas associated with each quantified metalloprotein and compared them to biogenic metal concentrations. From this, we estimated the proportion of total metal allocated to specific proteins and biogeochemically relevant metabolic pathways (see Methods). This approach provides a new framework for quantitatively linking metaproteomes with elemental function and stoichiometry in the ocean.

*Manganese allocation:* Under iron replete conditions, the majority of biogenic manganese was allocated to Mn-cluster proteins of the OEC (Fig. 4A), consistent with single cell SXRF measurements and theoretical predictions of metal use^14,54,67^. Notably, our estimates show Mn-cluster protein abundance often exceeded the amount of manganese available for their metalation, suggesting continued synthesis of these proteins under high iron despite insufficient manganese availability. Under low iron, 25-29% of total biogenic manganese was allocated to Mn-cluster proteins (Fig. 4A), despite increased manganese assimilation (Fig. 2E). This suggests that other manganese pools like storage^68^ and Mn-SOD^58,69^ may be important reservoirs of manganese under iron limitation, highlighting that the mechanistic basis for iron and manganese co-limitation requires consideration of additional cellular manganese pools beyond the OEC.

**Figure 4.**
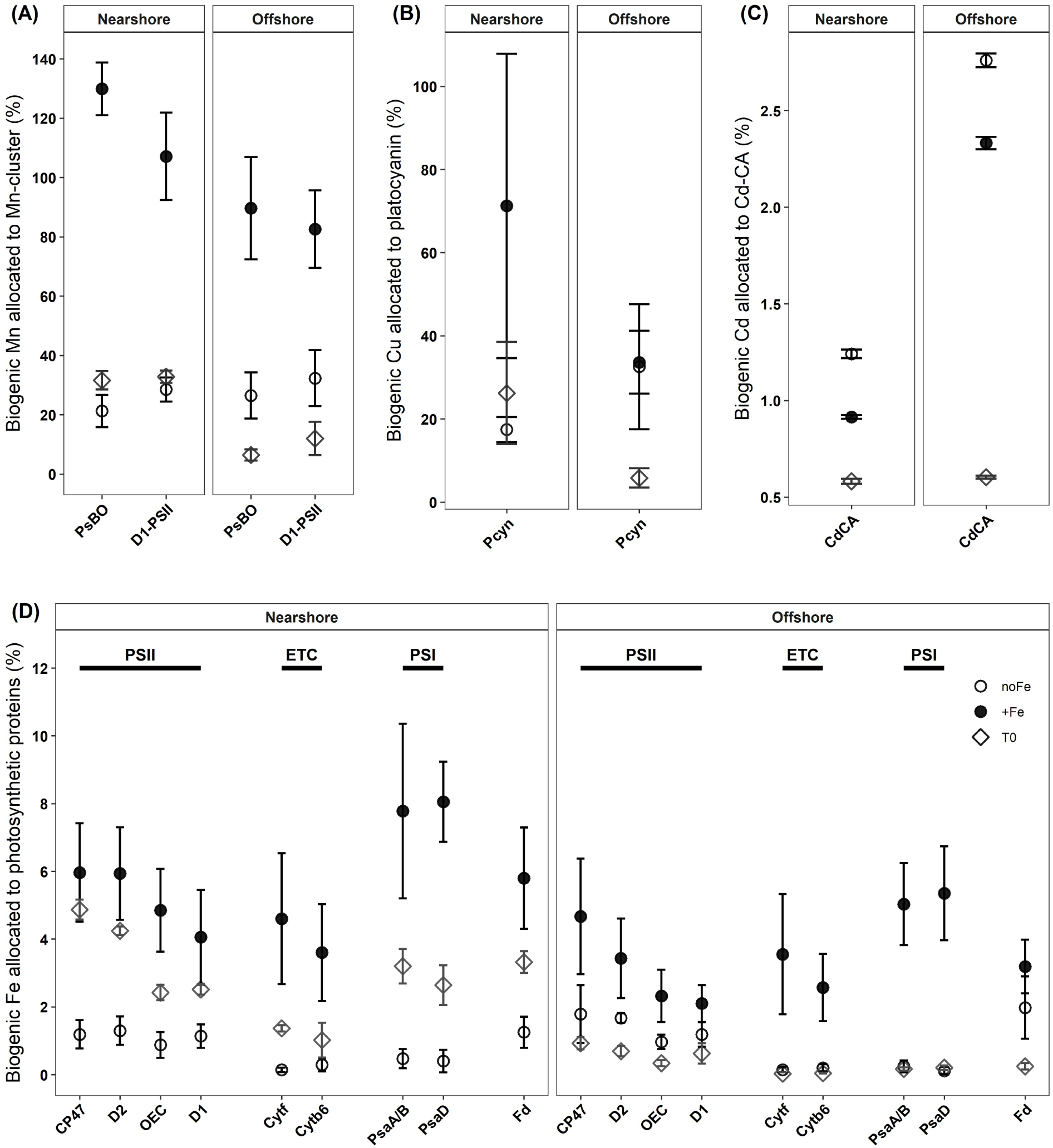
Trace metal allocations to metalloproteins containing manganese. **(A), copper (B), cadmium (C) and iron (D)**. In A-D points represent means of biological measurements and error bars represent *±* 1 SD. Diamonds represent in-situ (T0) samples, filled circles represent iron addition treatments, and open circles represent no iron added treatments. When possible, the averages of several proteins were used to calculate metal allocation to a specific protein complex (e.g. Mn cluster, PSII, ETC, PSI). In D) PSII = photosystem II, ETC = electron transport chain, PSI = photosystem I.

*Copper allocation*: Under iron replete conditions, 70% and 32% of biogenic copper was allocated to plastocyanin in the Nearshore and Offshore locations, respectively (Fig. 4B). This is consistent with previous estimates of copper allocation to plastocyanin in laboratory cultures^55,70^, and did not vary strongly with iron (Fig. 4B). The constitutive expression of plastocyanin and steady demand for plastocyanin-copper highlights the importance of this metal in seasonally iron-limited primary productivity in the SO.

*Cadmium allocation:* Overall, less than 3% of biogenic cadmium was allocated to Cd-CA, consistent with cadmium’s minimal biological role as a cofactor in this enzyme (Fig. 4C). We also observed increased cadmium assimilation under low iron (Fig. 2D), similar to prior observations^71,72^. This is most likely driven by non-specific uptake through upregulated iron transport pathways^73^, rather than a physiological response to low iron. Further examination of cadmium allocation within SO microbes would improve our understanding of how cadmium is managed or used within cells (e.g. are there safe storage strategies similar to those found in *Chlamydomonas*?^74^).

*Iron allocation:* Under iron replete conditions, 15–26% of biogenic iron was allocated to photosynthetic proteins (photosystem II, electron transport chain, photosystem I, and ferredoxin) (Fig. 4D), consistent with what has been reported in temperate diatoms^52^. Strikingly however, less than 5% of iron was allocated to these proteins under iron scarcity (Fig. 4D), contrasting previous estimates suggesting photosynthesis accounts for over half of cellular iron under low-iron conditions^52,54^. This result may be driven by methodological limitations. For example, it cannot be excluded that the strong down regulation of photosynthetic proteins under iron limitation (Fig. 2 G,H) led to an underestimation of their abundance as our proteomics methods may be biased against detecting low abundance proteins (see Methods). Possible background iron from sample processing and handling may also disproportionately affect estimates in low-iron treatments, potentially leading to artificially elevated bulk iron quotas and causing an underestimation of photosynthetic iron allocation. However, contamination during our experiments was thoroughly checked and found to be minimal. It is also possible that a fraction of the iron quota is reallocated to mitochondria for cellular maintenance under photosynthetically compromised conditions^75^. Indeed, we observed higher allocation of total protein to mitochondria under low iron compared to iron replete conditions (Fig. S3), consistent with previous reports of significant reduction in chloroplasts and increased mitochondrial proteins in iron-limited diatoms^75^.

### Decoupled ribosomal abundance and N:P stoichiometry

According to the growth rate hypothesis (GRH), faster organismal growth rates require more phosphorus-rich ribosomes to support protein synthesis, thereby increasing phosphorus demand and lowering cellular N:P^76,77^. In our study, iron enrichment increased ribosomal abundance within the phytoplankton community (Fig. 5A), linking increased protein synthesis (Fig. S4) and growth^33^ with increased ribosomal abundance. However, instead of lowering N:P, this increase in ribosomes was accompanied by increased N:P (Fig. 5B,C).

**Figure 5.**
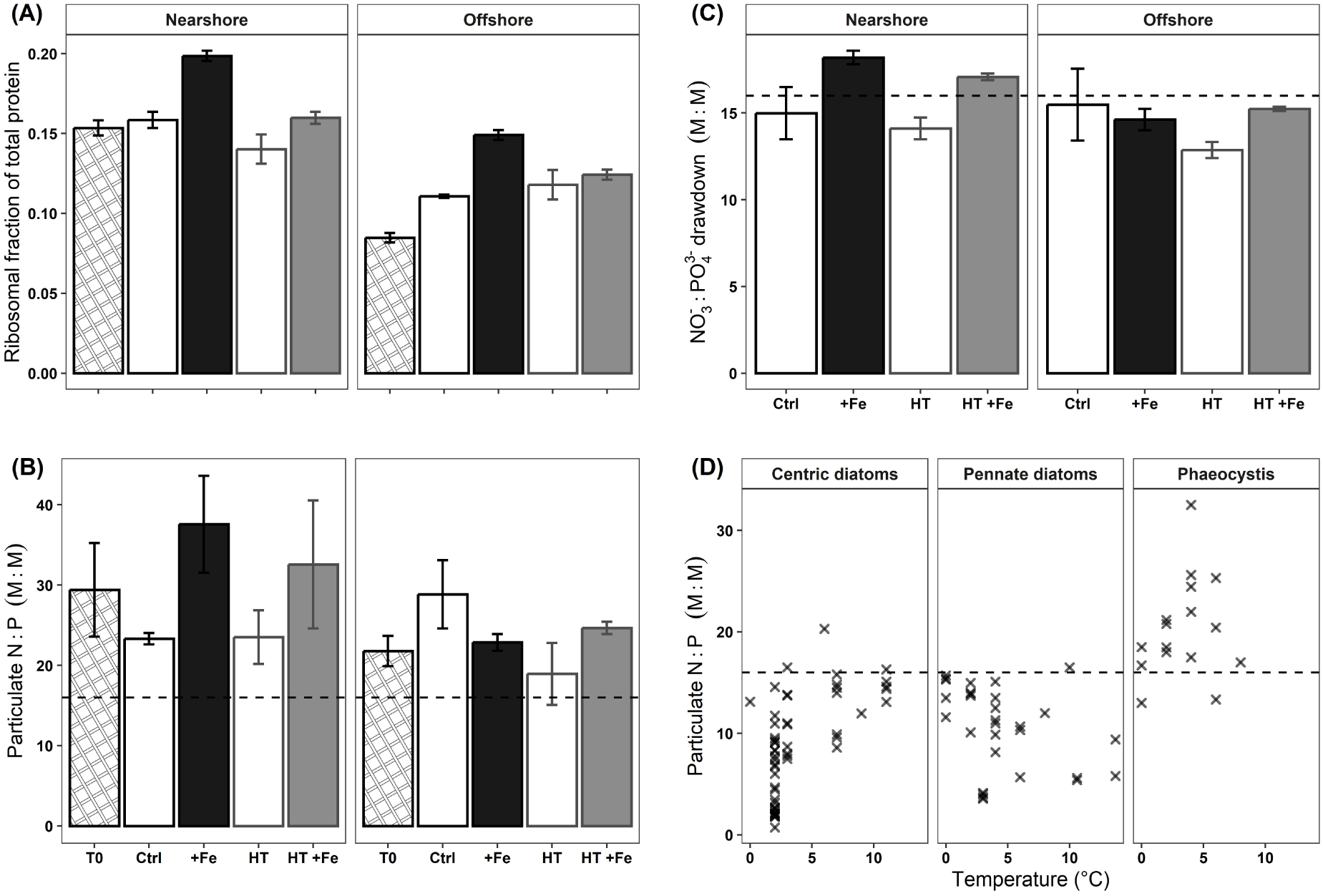
Ribosomal protein abundance increased with iron addition and decreased with warming, consistent with the growth rate hypothesis and translation compensation. However, ribosomal abundance was decoupled from N:P stoichiometry. **A)** Ribosomal protein fraction (sum of all ribosomal proteins) of total protein abundance (calculated as the sum of all matched peptide intensities) in-situ (T0) and after eight days of iron-temperature incubations. **B)** Ratios of bulk particulate nitrogen to particulate phosphorus (biogenic stoichiometry). **C)** Ratios of nitrate to phosphate drawdown (M:M), calculated using dissolved nitrate and phosphate measurements from day 2 and day 8 of iron-temperature incubations. **D)** Particulate (biogenic) N:P ratios acquired from previously published laboratory studies of cultures grown under nutrient replete conditions and various temperatures. Centric diatoms include *Thalassiosira* sp., *Chaetoceros* sp., *Skeletonema* sp., and *Odontella* sp. Pennate diatoms include *Pseudo-nitzschia* sp. and *Fragilariopsis* sp. See Fig. S5 for references. In A, B and C, bar plots represent means of triplicate measurements, and error bars represent ±1 SD and fall within the bounds of bars when not visible. In B, C, and D, dashed horizonal lines represent Redfield N:P ratio of N_16_:P_1_.

Conversely, warming improves translation efficiency per ribosome, thus fewer ribosomes are thought to be needed to sustain higher growth rates at higher temperatures, which raises N:P— a concept known as translation compensation^78–80^. Indeed, microbially derived particulate matter from warmer marine regions show higher N:P, and models project increased N:P in warming oceans^81–83^. Here we also observed decreased ribosomal abundance under warming (Fig. 5A), providing evidence for translation compensation in natural microbial assemblages. However, decreased ribosomal abundance under warming did not increase N:P (Fig. 5B,C) as predicted. This is consistent with other observations from the Ross Sea in the SO^43^, and suggests that factors beyond GRH or translation compensation influence the relationship between ribosomes, iron, temperature, and N:P in the SO.

First, increased nitrogen uptake after upregulation of iron-dependent nitrogen acquisition proteins under high iron (Fig. 2 G,H) can cause disproportionate accumulation of nitrogen relative to phosphorus investment in ribosomes, ultimately increasing N:P. Second, variations in organismal N:P^84,85^ and different stoichiometric responses to temperature can influence bulk N:P in complex phytoplankton communities, like those in our study (Fig. 5D). Third, intracellular nitrogen and phosphorus storage, especially in areas like the SO where both elements are available in high concentrations, could dampen changes in N:P due to fluctuations in ribosomal abundance^86^. Altogether, these findings suggest that multiple interacting factors likely drive N:P stoichiometric variability in the SO, with ribosomal abundance representing only one of them. This highlights the need for further research into the mechanisms shaping N:P in this region, and implications for macronutrient delivery to lower latitudes.

## Conclusions

By integrating metaproteomics with elemental measurements, we establish a mechanistic framework for elemental allocation to key metabolic processes within natural microbial assemblages. Using this approach, we observed high metabolic flexibility within SO phytoplankton, likely contributing to their ecological success in this dynamic environment. Specifically, we show tunable PSI:PSII architecture, compensatory use of non-iron metals, and reallocation of elements to various metabolic processes in response to shifting physiological requirements. Furthermore, we quantitatively constrain a proportion of biogenic metal to various proteins, providing insights into the molecular mechanisms driving elemental demand and limitation. Lastly, our findings suggest that factors beyond ribosomal abundance, such as storage, play a substantial role in influencing N:P stoichiometry, and should be considered when examining macronutrient distribution in the SO and beyond.

Looking ahead, the use of next-generation mass spectrometers^87^ alongside paired nucleotide databases will enable the generation of deeper metaproteome profiles within shorter timeframes. While this will necessitate addressing novel computational challenges associated with the scale and complexity of the resulting data, it will profoundly accelerate our ability to measure elemental allocation across a wider range of proteins and biogeochemical processes, including at taxon-specific resolutions. These advances could also facilitate the discovery of molecular indicators of intracellular storage, which is increasingly recognized to be essential for understanding marine elemental dynamics.

In sum, our work bridges molecular data with biogeochemical processes, and provides a path towards incorporating omics-based measurements into biogeochemical models that rely on elemental stoichiometry in their parameterizations.

## Methods

### Experimental setup

We conducted two shipboard iron-temperature experimental manipulations during the austral summer of 2018-2019 in the Weddell Sea, Southern Ocean^33^. The first experiment (Offshore) was initiated on December 28, 2018, approximately 560 km away from the ice edge at 65 °S 0.038 °E. The second experiment (Nearshore) was initiated on January 9, 2019, approximately 70 km from the ice edge at 70.15 °S 11.02 °W (Fig. 1A). For both experiments, we collected sea water at ∼21 m depth using a custom built ultraclean CTD sampling system – “Titan” – equipped with PRISTINE high-volume samplers^34,35^ made from polypropylene to be light proof rather than the original PVDF version. This depth minimized trace metal contamination from the ship and coincided with phytoplankton biomass maxima (Fig. S1). Titan was moved into a dedicated, temperature-controlled, trace-metal-clean (TMC) laboratory container where sea water was used for initial in-situ (T0) sampling and to fill triplicate, 20 L, TMC cubitainers for each iron-temperature treatment.

The treatment matrix included four iron-temperature conditions: *i*) a control at in-situ temperature and no iron addition (*Ctrl*), *ii*) iron (^57^Fe) addition at in-situ temperature (*+Fe*), *iii*) warming—high temperature—with no iron addition (*HT*), *iv*) concurrent warming and iron (^57^Fe) addition (*HT +Fe*). We supplemented the high iron treatments with a naturally occurring but very rare isotope of ^57^Fe, which allowed us to differentiate between the added iron and iron already present in the water (predominantly ^56^Fe). For this, we made a 40 μM ^57^FeCl_3_ stock solution in 0.1 M trace-metal grade HCl, then added 1 mL of this stock to each of the high iron cubitainers. This resulted in a total initial dissolved ^57^Fe concentration of 2 nM. We then sealed the cubitainers and placed them inside temperature-controlled deck incubators where the low temperature treatments were incubated under in-situ temperature, and the high temperature treatments were incubated under in-situ +2 °C (Table 1). We note that the incubators could not reliably maintain the frigid in-situ temperature of the Nearshore location, so we adjusted the temperature by an additional 0.4 °C for all treatments in this experiment. We used black mesh screens to keep light at 15-20% of incident inside the incubators, which ensured light-saturated growth without inducing photodamage. We maintained the incubations for eight days. Throughout the experiments, we followed previously described TMC protocols for cleaning, water collection, and sample processing^88–90^.

### Macronutrient measurements

*Dissolved macronutrients –* We measured nitrate, phosphate and silicate (silicic acid) on board the ship using a four channel Auto-Analyzer (QuAATRo, Seal Analytical, Germany), following the Joint Global Ocean Flux Study protocols described in (Gordon et al., 1993). Briefly, we syringe filtered 125 mL from each biological replicate through 0.2 μm Acrodisc® filters, and collected the filtrate in polyethylene bottles that were rinsed three times with the filtrate before use. When immediate measurements were not possible, we stored the samples in capped bottles at 4 °C for a maximum of five hours. We made ten-point calibration curves daily by diluting stock solutions of the different nutrients in 0.2 μm filtered low-nutrient seawater. We also analyzed a certified reference material (KANSO, Japan) containing known concentrations of nitrate, phosphate, and silicate in Pacific Ocean water with each run. All samples and standards were brought up to 21 °C before measurements were taken.

*Particulate macronutrients –* We collected samples for particulate organic carbon (POC) and particulate nitrogen (PN) using vacuum filtration through pre-combusted GF75 glass microfiber filters (Whatman, 0.3 μm pore size, 25 mm diameter). We filtered 1 L from each biological replicate, wrapped the filters in aluminum foil, and then stored them at −20 °C until further laboratory analysis. We analyzed POC and PN using Thermo-Interscience Flash EA1112 Series Elemental Analyzer (Thermo Scientific) with a detection limit of 100 ppm and a precision of 0.3 % ^91^. Briefly, samples were flash combusted with excess oxygen at 900 °C to convert organic carbon and particulate nitrogen into CO_2_ and N_2_ gas, respectively. A helium carrier gas was then used to transfer CO_2_ and N_2_ into the elemental analyzer for measurement. We also measured filter and sample handling blanks, and subtracted the blank measurements from sample measurements. The sampling and analysis methods for particulate phosphorus are described in the trace metal measurements section below.

### Trace metal measurements

We took samples for dissolved and particulate trace metals, and performed all the metal (and particulate phosphorus) measurements using High Resolution Inductively Coupled Plasma Mass Spectrometry (HR-ICP-MS). For this, we followed strict TMC protocols during sampling and sample processing^88–90^.

*Dissolved trace metal sampling and sample processing* – To separate the dissolved trace metal fraction from the particulate fraction, we filtered water from each biological replicate through pre-cleaned 0.2 μm filter cartridges (AcroPak). We collected the filtrate in acid cleaned 125 mL LDPE bottles that were rinsed five times with the filtrate before use. We then acidified the samples using trace-metal grade Baseline^®^ HCl (Seastar Chemicals Inc) to a final concentration of 0.024 M HCl, which resulted in a pH of ∼1.8. We stored the samples at room temperature until further laboratory analyses. In the laboratory, we preconcentrated the dissolved trace metals from the acidified samples following methods in^92^. Briefly, we passed each sample through a SeaFAST-pico preconcentration system (ESI, USA), with two 10 mL loops for a total sample volume of 20 mL. With this, we concentrated each sample into 350 µL of elution acid (double distilled 1.5 M HNO_3_) containing rhodium (Rh) as internal standard, and then stored at room temperature until further analysis with HR-ICP-MS.

*Particulate trace metal and phosphorus samples* – We used polyethersulfone (PES) filters (25 mm, 0.45 μm Pall Supor) placed on polypropylene filter holders (Advantec) with polypropylene Luer Locks (Cole-Palmer) to capture the particulate trace metal fraction. We pre-cleaned the filters by soaking in distilled 1.2 M HCl (VWR Chemicals – AnalaR NORMAPUR) at 60 °C for 24 h, then rinsing with ultrapure MQ water 5 times, and eventually storing in ultrapure MQ water until use^93^. After sampling, we stored the PES filters in acid-cleaned Eppendorf vials at −20 °C until further laboratory analysis.

We digested the particulate trace metal samples in two successive steps to capture the labile and refractory fractions separately^89,90^. We summed the measurements from these two fractions to get the total particulate metal and phosphorus measurements shown in this study. Briefly, we first solubilized the labile fraction by subjecting the PES filters to a weak leach, consisting of 1.8 mL solution of 4.35 M acetic acid (double distilled using PFA Savillex stills, VWR NORMAPUR) and 0.02 M hydroxylamine hydrochloride (Sigma-Aldrich – 99.999% trace metal basis). The vials were heated to 95 °C for 10 minutes in a water bath and were subsequently cooled down to room temperature. After two hours, the filters were moved to a 30 mL Teflon vial (Savillex), and the remaining leachate was centrifuged at 16000 RCF for 10 minutes to bring any remaining particles to the bottom of the solution in the vials. Next, 1.6 mL of leachate was transferred to a separate 30 mL Teflon vial (Savillex) and 100 μL of concentrated HNO_3_ (double distilled, AnalaR NORMAPUR) was added. The vials were then heated to near dryness at 110 °C, re-dissolved in 2 mL of 1.5 M HNO_3_ with 10 ng mL^-1^ Rh as internal standard, and stored at room temperature for subsequent analysis by HR-ICP-MS.

For the refractory fraction, we transferred the remaining 0.2 mL of leachate from the labile leach vials to the Teflon vials already containing the PES filters – we corrected for the 0.2 mL labile leachate in the refractory fraction. Next, we added a 2 mL solution of 8.0 M HNO_3_ (triple distilled, VWR Chemicals – AnalaR NORMAPUR) and 2.9 M HF (Supelco, Ultrapur), and refluxed the capped samples for 4 hours at 110 °C. Afterwards, we cooled the samples to room temperature and poured the contents into secondary Teflon vials (Savillex) without transferring the filters. We then heated the secondary Teflon vials to near dryness at 110 °C, redissolved the contents in a 1 mL a solution of 8.0 M HNO_3_ and 15% H_2_O_2_ (Merck – Suprapur), refluxed for second time for 1 hour at 110 °C, and again heated to dryness at 110 °C. Lastly, we redissolved the samples in 2 mL of 1.5 M HNO_3_ with 10 ng mL^-1^ Rh as internal standard, and stored them at room temperature for subsequent analysis by HR-ICP-MS.

*HR-ICP-MS methods* – We analyzed the samples using Thermo Scientific Sector Field High Resolution Element 2 mass spectrometer, with a microFAST introduction system (ESI, USA) equipped with a PFA nebulizer. Detailed HR-ICP-MS methods are described in^94^. We blank-corrected the measurements for contaminants from sample handling, reagents and filters, following^89,90^. The accuracy and precision of the measurements were determined by measuring GEOTRACES community consensus reference materials SaFe D1, GSC and GSP as well as in-house reference seawater samples. Rhodium was used as an internal standard for particulate samples and indium and lutetium for dissolved samples, and drift standards were measured after every batch of 13 samples to correct for drift during the runs. We calculated trace metal and phosphorus concentrations using a mixed solution of elemental standards.

### Metaproteome sampling and sample preparation

We used a peristaltic pump system to sequentially filter 2-4 L from each biological replicate through 12, 3.0, and 0.2 µm polycarbonate filters (Isopore, 47 mm diameter). Filters were then stored at −80 °C until further laboratory analysis.

We extracted and digested proteins from filters using methods previously described in^95^. Briefly, 600 µL of sodium dodecyl sulphate (SDS) lysis buffer (2% SDS, 0.1 M Tris/HCl pH 7.5, 5% glycerol, 5 mM EDTA) was added to each sample, followed by a 10-minute incubation on ice then a 15-minute incubation in a benchtop thermomixer (Eppendorf Thermomixer C) at 95°C and 350 RPM. Samples were then sonicated on ice for one minute using a QSonica125 microprobe (50 % amplitude, 125 W, 15 second on/off pulses) and left at room temperature for an additional 30 minutes. The extracted proteins, which were suspended in SDS lysis buffer at this point, were purified by removing the filters and centrifuging the samples at 15000 ×g for 30 minutes to pellet cellular debris. After this, we acquired the protein-containing supernatant and pooled the three different size fractions from each biological replicate by combining an equal volume from each fraction. We quantified bulk protein concentrations using a Pierce Micro BCA Protein Assay Kit following manufacturer’s protocol.

We used S-Trap mini columns (PROTIFI) to remove SDS lysis buffer from the samples and digest the extracted proteins. Samples were first reduced and alkylated using 5 mmol L^-1^ dithiothreitol and 15 mmol L^-1^ iodoacetamide respectively, then denatured using 12% phosphoric acid. Samples were then diluted with S-trap buffer (pH 7.1, 100 mmol L^-1^ tetraethylammonium bicarbonate in 90% aqueous methanol) (1sample:7buffer, v:v) and loaded onto the S-trap columns. Once on column, we washed each sample 10 times with 600 µL S-trap buffer to remove the SDS. Proteins were then digested on-column using trypsin (1trypsin:25protein, µg:µg) at 37 °C for 16 hours. Tryptic peptides were eluted from the S-Trap columns with a series of 50 mmol L^-1^ ammonium bicarbonate, 0.2% aqueous formic acid, and 50% acetonitrile 0.2% formic acid washes. Samples were then desalted on pre-conditioned 50 mg C18 columns (HyperSep), dried using a Vacufuge Plus (Eppendorf), and finally resuspended in a 1% formic acid 3% acetonitrile solution prior to liquid chromatography – tandem mass spectrometry (LC-MS-MS).

### LC-MS-MS

We performed duplicate 1 µg injections for each biological replicate on a Dionex UltiMate 3000 RSLC-nano LC system (Thermo Scientific) coupled to a Q-Exactive hybrid quadrupole-Orbitrap mass spectrometer (Thermo Scientific). Peptides were separated by reverse phase chromatography under a non-linear gradient (Table S1) with a self-packed column (50 cm x 0.1 mm ID) containing JUPITER Proteo C18 resin. The mass spectrometer was operated in data-dependent acquisition mode – detailed mass spectrometer settings are shown in (Table S2).

### Bioinformatics and data analysis

We assessed instrument and acquisition performance by analyzing the unprocessed raw mass spectrometry files using RawTools v2.0.7^96^. We then converted the – quality checked – raw files into standardized mzML format using ThermoRawFileParser v1.4.0^97^ before further analysis. *Reference database –* We used a reference SO metatranscriptome database^43^ to match the acquired mass spectra with peptides. Briefly, samples for this database were collected from surface water in the McMurdo Sound, SO, on January 15^th^, 2015 which were subjected to experimental incubation as described in^43^. Sequencing was performed via Illumina HiSeq, transcript contigs were assembled de novo using CLC Assembly Cell, and open reading frames (ORFs) were predicted using FragGeneScan^98^. We filtered ORFs to eliminate those with less than 50 reads total across all database samples, and annotated for putative function using hidden Markov models and BLAST-p against PhyloDB^99^. ORFs were assigned a taxonomic affiliation based on best Lineage Probability Index^99,100^. Lastly, we grouped ORFs into protein clusters by sequence similarity using Markov clustering^101^, and consensus cluster annotations were based on KEGG, KO, KOG, KOG class, Pfam, TIGRfam, EC, GO.

*Database searching –* We matched the acquired mass spectra with peptide sequences from the reference database using MSGF+ within OpenMS^102,103^. We first appended the reference database with a database of common contaminants from the common Repository of Adventitious Proteins (The Global Proteome Machine Organization) and removed redundant sequences using Python script by P. Wilmarth (github.com/pwilmart/fasta_utilities). We conducted the search using a 10-ppm precursor mass tolerance, and set cysteine carbamidomethylation as a fixed modification, and methionine oxidation and N-terminal glutamate to pyroglutamate conversion as variable modifications. Lastly, we filtered peptide-spectral matches at a 1% false discovery rate using a target-decoy strategy, where ‘target’ represents all the sequences in the database and ‘decoy’ represents all the database sequences in reverse^104^.

*Peptide quantification –* We grouped the duplicate LC-MS-MS injections for each biological replicate and aligned the retention times within each of these groups using MapAlignerIdentification^105^. We then matched unknown features in one injection with identified peptide features from the other injection, which allowed us to improve the number of peptide identifications in each sample. We then calculated feature intensity (i.e. relative peptide abundance) based on MS1 ion intensity using FeatureFinderIdentification^106^.

*Peptide annotation –* We mapped peptides to corresponding ORFs for protein cluster assignment, and for functional and taxonomic annotations. Peptides mapping to one or more ORFs from the same cluster or functional annotation were annotated with that cluster / function. Peptides mapping to ORFs with different clusters / functional annotations were annotated as “ambiguous cluster” / “ambiguous function”. We then assigned a taxonomy string to each peptide by resolving taxonomic affiliation at the genus level, or, when not possible, by gradually progressing through taxonomic rank until a non-ambiguous taxonomic level is found. That is, peptides mapping to one or more ORFs from the same genus (e.g., *Fragilariopsis*) were annotated down to genus (eukaryote-stramenopile-bacillariophyte-pennate-Fragilariopsis). Peptides that were ambiguous at the genus level (e.g., mapping to both Fragilariopsis (pennate diatom) and Chaetoceros (centric diatom) ORFs) were annotated down to the class level as “ambiguous diatom”, and so on. Peptides that could not be resolved at the domain level were annotated as “ambiguous domain”.

*Database suitability and normalization–* Untargeted metaproteomics experiments like ours generate relative abundance measurements that must be normalized before making comparisons across samples. Several factors influence these abundance values and complicate their normalization, including the suitability of the reference database and heterogeneity of non-peptide features across samples. Here, we leveraged previously established methods in^107^ to test database suitability and to find an appropriate normalization method for our data.

We calculated and correlated three different abundance metrics using our metaproteomics data. I) We calculated the summed total ion current (TIC) for each sample using pyOpenMS. TIC is a database independent sum of all mass spectral peak intensities measured throughout a run, and therefore includes background noise and contamination, in addition to all measured peptides. II) We identified peptide-like features based on feature resemblance to the isotopic distribution of a theoretical peptide model, “averagine”^108^, and quantified their total abundance for each sample using Hardklör and Krönik^109^. The total abundance of peptide-like features is also database independent but is less influenced by background noise and contamination. III) We calculated the abundance of all matched peptides for each sample (see previous methods on database searching and peptide quantification). This approach is database dependent and can therefore be biased by database suitability. A strong correlation between these three metrics would indicate that either one can be used for normalizing our relative abundance measurements (Fig. S6).

*Community composition –* We calculated the fractional contribution of various taxonomic groups to the metaproteome by summing the abundance of peptides within each taxonomic group and dividing by the total abundance of all matched peptides (e.g., sum of all centric diatom peptide intensities / sum of all peptide intensities). We focused our analyses on taxonomic resolutions with a high enough number of identified peptides for confident comparison of protein expression across and within treatments (Fig. S7). These taxa were diatoms and *Phaeocystis*.

*Proteins and protein group abundances –* We first used a coarse-grained clustering approach to pool peptides into four distinct metabolic categories: light-reaction photosynthetic proteins, RuBisCO, ribosomal proteins, and mitochondria. For this, we searched protein annotations against character strings that capture these groups. The photosynthesis group included all peptides with ‘photosyn*’ or ‘light-harvesting complex’ or ‘flavodoxin’ or ‘plastocyanin’ in their annotation. The RuBisCO group included all peptides with ‘RuBisCO’ or ‘rubisco’ or ‘bisphosphate carboxylase’ in their annotation. The ribosome group included all peptides with ‘ribosom*’ in their annotation. The asterisk (*) in the character string represents a wildcard character. We also used a finer grained approach to examine the abundance and differential expression patterns of specific proteins (e.g. Flavodoxin). For this, we searched protein annotations against character strings specific to the proteins of interest and used MCL clusters to define a protein/protein group.

*Quantitative trace metal allocation –* We first converted the relative abundances of each metalloprotein (unitless) into protein mass concentrations (ug L^-1^) (eq. 1). To do this, we normalized the summed peak areas of metalloprotein-specific peptides by the total abundance of peptide-like features in each sample (both unitless) (Fig. S8). Peptide-like feature abundance was quantified using Hardklör and Krönik^109^ and was assumed to be proportional to the total protein concentration loaded onto the mass spectrometer. This approach is database independent and is more resilient against missing peptide matches arising from database mismatches, posttranslational modifications, or other stochastic sources of error.

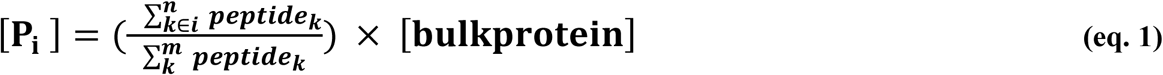

Where:

**[P_i_]** = mass concentration of metalloprotein of interest (P_i_), μg L^-1^

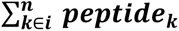 = summed intensities of all peptides uniquely matching the metalloprotein of interest (P_i_), unitless

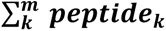 = summed intensities all peptide-like features (includes matched and unmatched features), unitless

**[bulkprotein]** = bulk protein concentration in sea water, μg L^-1^

Next, we converted the mass concentration of each metalloprotein (P_i_) into a molar concentration (i.e. copy numbers) using molar masses from previously published amino acid sequences (eq. 2).

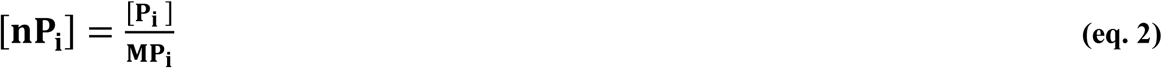

Where:

**[nP_i_]** = molar concentration of metalloprotein P_i_, mol L^-1^

**[P_i_]** = mass concentration of protein of interest (P_i_) calculated in eq.1, μg L^-1^

**MP_i_**= molar mass of metalloprotein P_i_ derived from genome sequences and previous studies, μg mol^-1^

We then calculated the equivalent molar concentration of metal that would be co-factored by each metalloprotein (eq. 3). The metal-to-protein molar ratios were obtained from previous studies, and for simplicity, we assumed perfect metal-protein specificity. That is, each protein binds exclusively to its known metal cofactor. This parameterization may, however, be prone to error due to variability and uncertainty in protein-metal binding, and especially under varying environmental conditions^11,110^. To account for the uncertainty in metalation efficiency, we introduced a metalation coefficient, σ, randomly sampled between 0.5 (i.e., 50% of measured metalloproteins are metalated) and 1 (i.e., complete metalation). This calculation was repeated 1,000 times to incorporate variability.

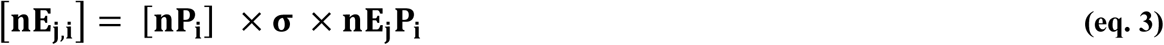

Where:

**[nE_i,j_]** = molar concentration of metalloprotein (P_i_)-associated metal (Ej), mol L^-1^

**[nP_i_]** = molar concentration metalloprotein P_i_ calculated in eq. 2, mol L^-1^

**σ** = metalation coefficient, where 1 = perfect metalation (i.e. every metalloprotein molecule is bound to its metal cofactor), 0 = no metalation (i.e. no metalloprotein molecule is bound to its metal cofactor), unitless

**nE_j_P_i_** = moles of metal (E_j_) per moles of protein (P_i_), e.g. 1 = one metal atom per protein, unitless

Lastly, we determined the contribution of each metalloprotein to the total biogenic metal reservoir, expressed as a fraction of the particulate metal pool measured by ICP-MS (eq. 4)

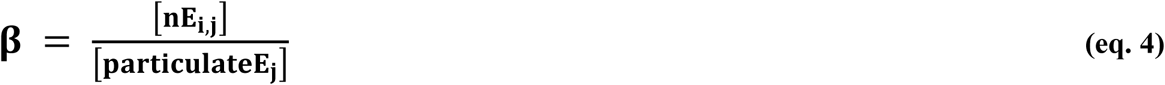

Where:

**β** = fraction of total biogenic metal found in metalloprotein of interest (Pi), unitless

**[nE_i,j_]** = molar concentration of metalloprotein (P_i_)-associated metal (E_j_) as calculated in eq. 3, mol L^-1^

**[particulateE_j_]** = molar concentration of particulate (biogenic) metal measured using ICP-MS, mol L^-1^

All together:

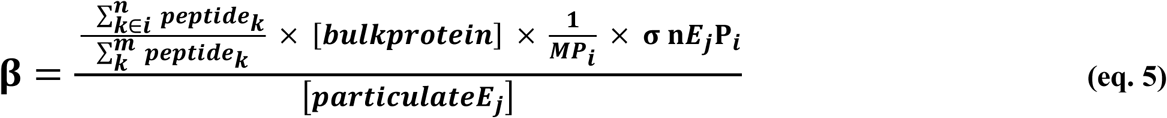

### Fragilariopsis cylindrus cultures and proteomics

We grew *Fragilariopsis cylindrus* (NCMA 1102) in EDTA buffered Aquil* media under trace-metal clean conditions (as described in^32,63^) and the following treatments: 1) low iron (low Fe: 15 nM Fe + 48 nM Mn); 2) medium iron (med Fe: 35 nM Fe + 48 nM Mn); 3) low manganese (low Mn: 500 nM Fe + no added Mn); 4) replete (replete: 500 nM Fe + 48 nM Mn). Temperature was maintained at 3 °C, and constant light was supplied at ∼ 50 μmol of photosynthetically active radiation (PAR) m^−2^ s^−1^. Cells were filtered on 0.2 μm polycarbonate filters prior to sample processing and LC-MS-MS measurements^32^.

### Statistical analysis and visualization

We performed all statistical analyses and visualizations using various packages in R v4.2.1. Differential expression and fold-change analyses were conducted on taxon normalized protein cluster abundance values using Bayes quasi-likelihood F-tests with the glmQLFTest function in edgeR v4.2^111^. For this, we first filtered that data to remove any protein clusters with fewer than two matched peptides. We then imputed missing values via random forest-based imputation^112^ using the missForest v1.5 package^113^, and excluded any proteins that were not detected in at least 75% of all samples. An FDR adjusted P-value cutoff of 0.05 was used for statistical significance. NMDS ordination analysis was conducted in two dimensions with 1000 maximum random starts using the Vegan v2.6.2 package. The R packages ggplot2 v3.4.1 networkD3 v0.4, cowplot v1.1, and ggOceanMaps v1.2.6 were used for data visualization.

## Data and code availability

All proteomics data including the raw mass spectrometry files and processed data files have been deposited to the ProteomeXchange Consortium via the PRIDE partner repository^114^ with the dataset identifier <u>PXD042627</u> and <u>10.6019/PXD042627</u>; reviewer account details - username, password: eabSxs2x. All other metadata are available at https://doi.org/10.25850/nioz/7b.b.hh. All code for bioinformatics analyses, statistical testing, and data visualization is available at: https://github.com/LoayJabre/PS117_metaP

## Acknowledgments

We thank RV Polarstern captain and crew, Dr. Olaf Boebel, Chief Scientist for PS117 expedition, and the PS117 science team. We are indebted to Sharyn Ossebaar for taking the dissolved macronutrient measurements onboard the ship, and to John Seccombe from Aquahort for his support in conceiving and building the incubators and for his remote support during the expedition. We thank Hung-An Tian, Indah Ardiningsih and Sven Pont for help with running the onboard experiments. We thank Patrick Laan for help with taking the trace metal measurements. We thank Scott McCain for meaningful discussion and feedback. This study was funded by NSERC Discovery Grant RGPIN-2015-05009 to E.M.B, Simons Foundation Grant 504183 to E.M.B, Simons Foundation CBIOMES Award ID 1001702 to E.M.B, Canada Research Chair Support to E.M.B; ALWPP.2016.020 from the Dutch Research Council (NWO) to R.M, and the Nova Scotia Graduate Scholarship to L.J.J. The International GEOTRACES Programme is possible in part thanks to the support from the U.S. National Science Foundation (Grant OCE2123354 to TMC) to the Scientific Committee on Oceanic Research (SCOR).

## Author contributions

L.J.J performed data analyses and wrote the manuscript with input from the other authors. L.J.J performed laboratory culturing experiments, processed the metaproteomics samples, and analyzed the metaproteomics data under the supervision of E.M.B. E.R processed the proteomics samples from the cultures and performed the mass-spectrometry measurements. E.M.B, R.M, and C.P.D.B designed the incubation experiments with input from authors. M.M, C.E, E.M.B, and R.M collected all the samples aboard RV Polarstern. M.M took the elemental measurements. E.M.B, R.M and C.P.D.B supervised the project and acquired the funding.

