## Supporting Information for "Elemental allocation to molecular drivers of biogeochemistry in the Southern Ocean"

**ORCIDs**

**Loay J. Jabre:** 0000-0003-3177-6824

**Elden Rowland:** 0000-0003-4756-9125

**Charlotte Eich:** 0009-0009-3793-7896

**Mathijs van Manen:** 0000-0002-6016-916

**Corina Brussaard:** 0000-0002-6320-9229

**Rob Middag:** 0000-0002-3326-530X

**Erin M. Bertrand:** 0000-0002-5950-6810

**This document includes:**

- Figures S1-8
- Tables S1-S2

**Supplemental Figures**

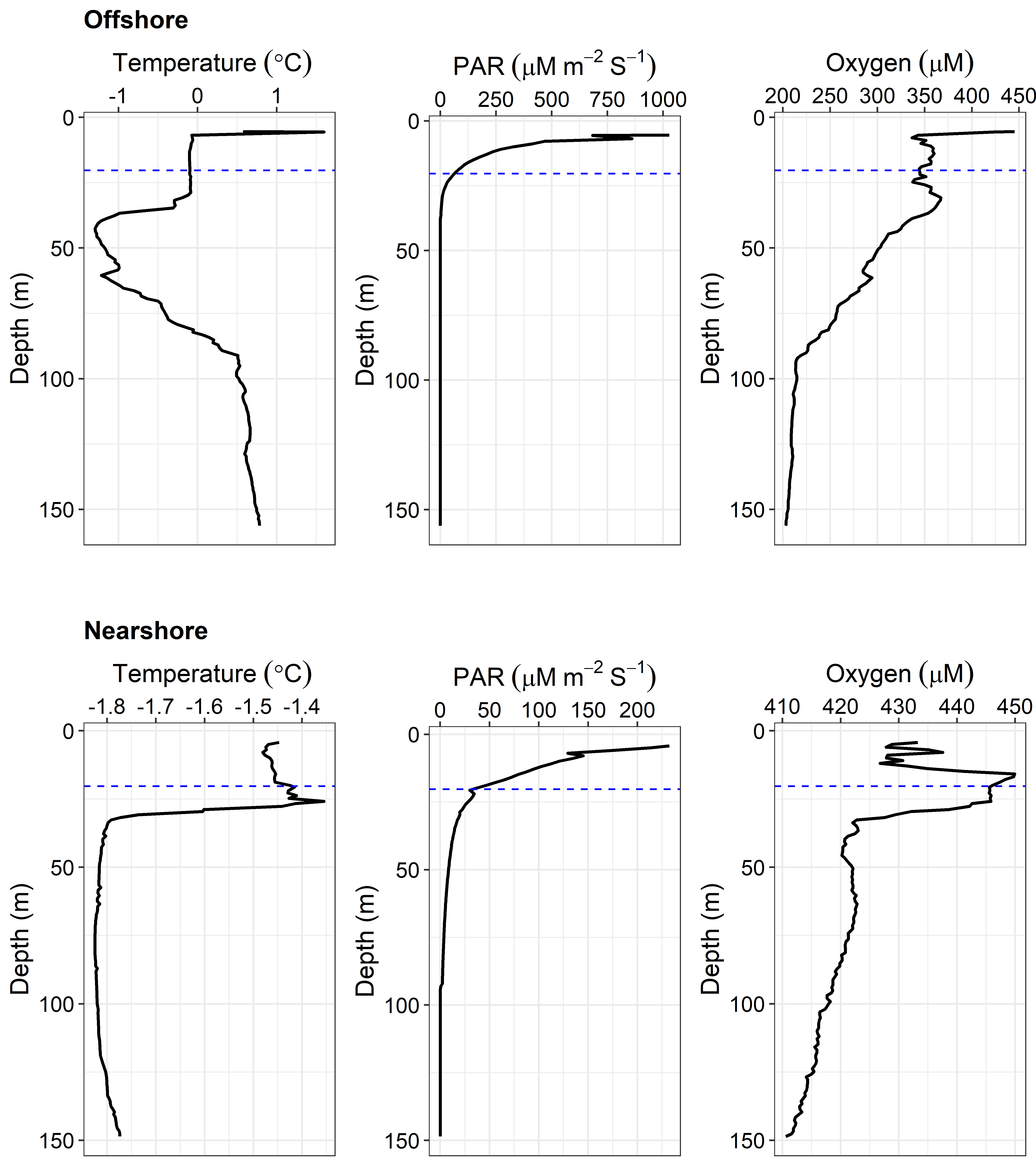

**Figure S1 –** Temperature, photosynthetically active radiation (PAR) and oxygen concentrations from the CTD downcast prior to water collection for the bioassay incubations. Horizontal dashed lines indicate sampling depth at each location. Due to the lack of chlorophyll-a sensor data on these casts, we used oxygen measurements as a proxy for photosynthesis to determine phytoplankton biomass maxima.

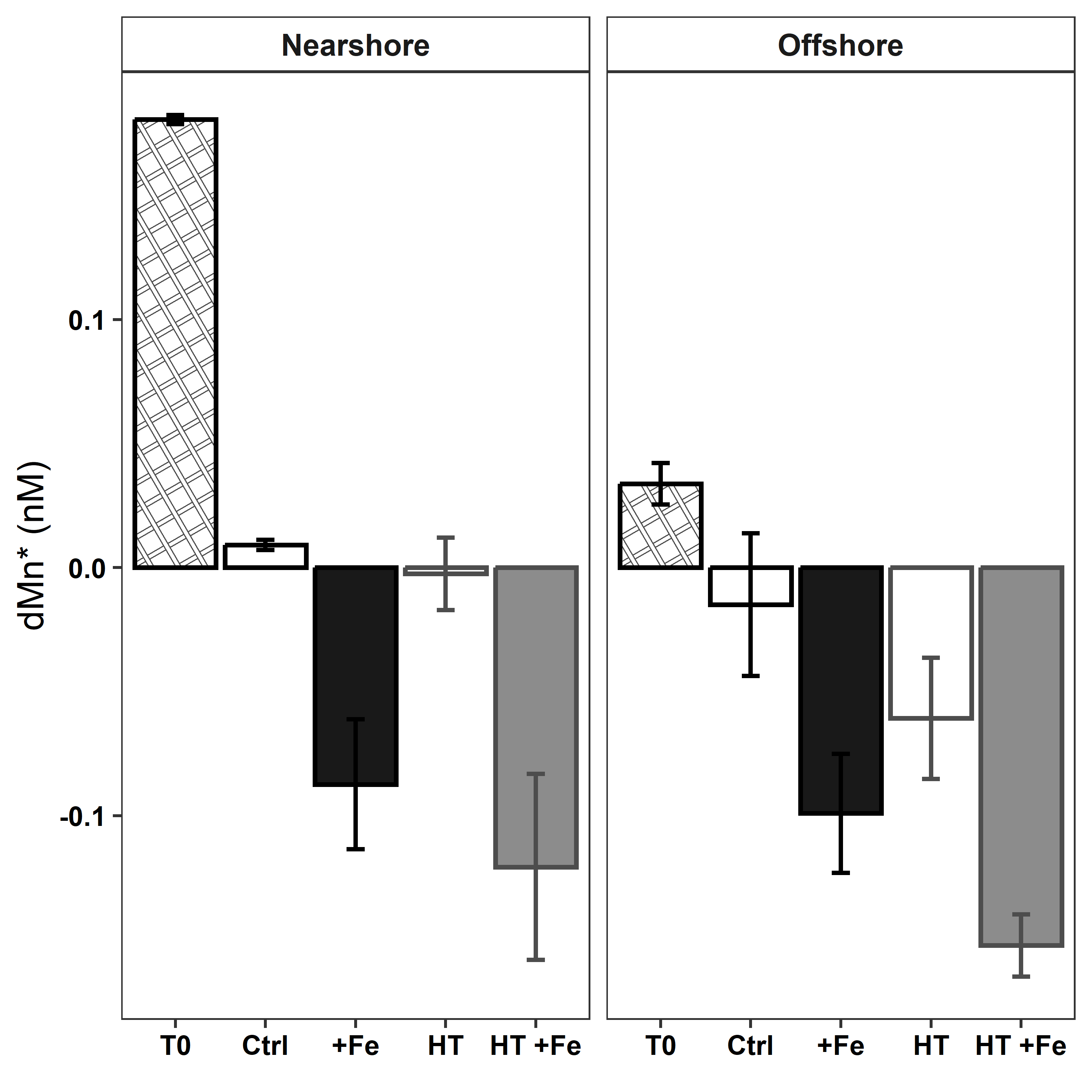

**Figure S2 –** Mn* in-situ (T0), and after eight days of incubation with and without iron addition. Mn* was calculated following^1^ as Mn* = [dMn] – [d^56^Fe56+d^57^Fe]/2.67.  Error bars represent ±1 SD. Lower Mn* values indicate stronger manganese limitation.

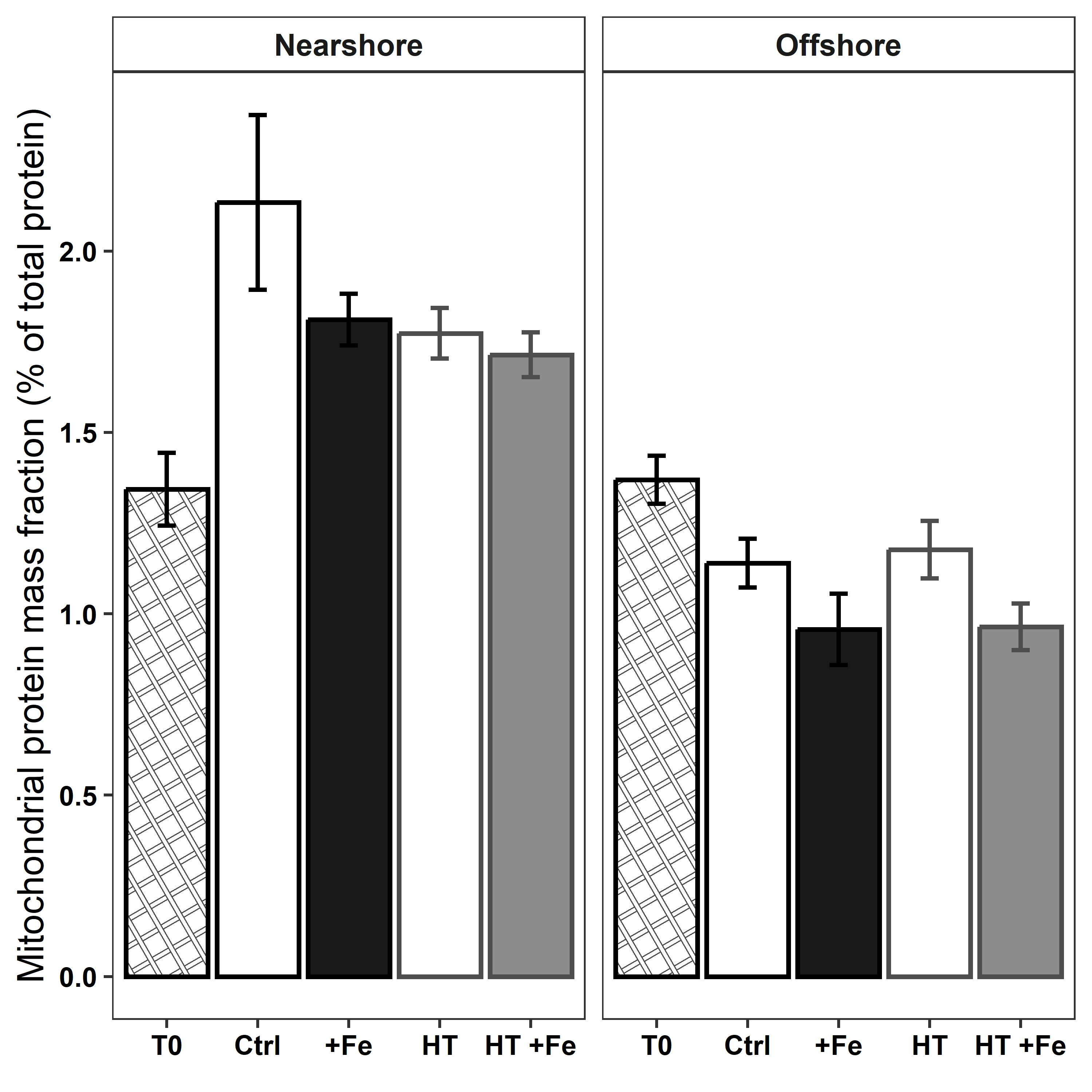

**Figure S3 –** Mitochondrial protein mass fraction in-situ (T0), and after eight days of incubation with and without iron addition. Mass fraction was calculated as the summed abundance of all mitochondrial peptides divided by total peptide abundance.

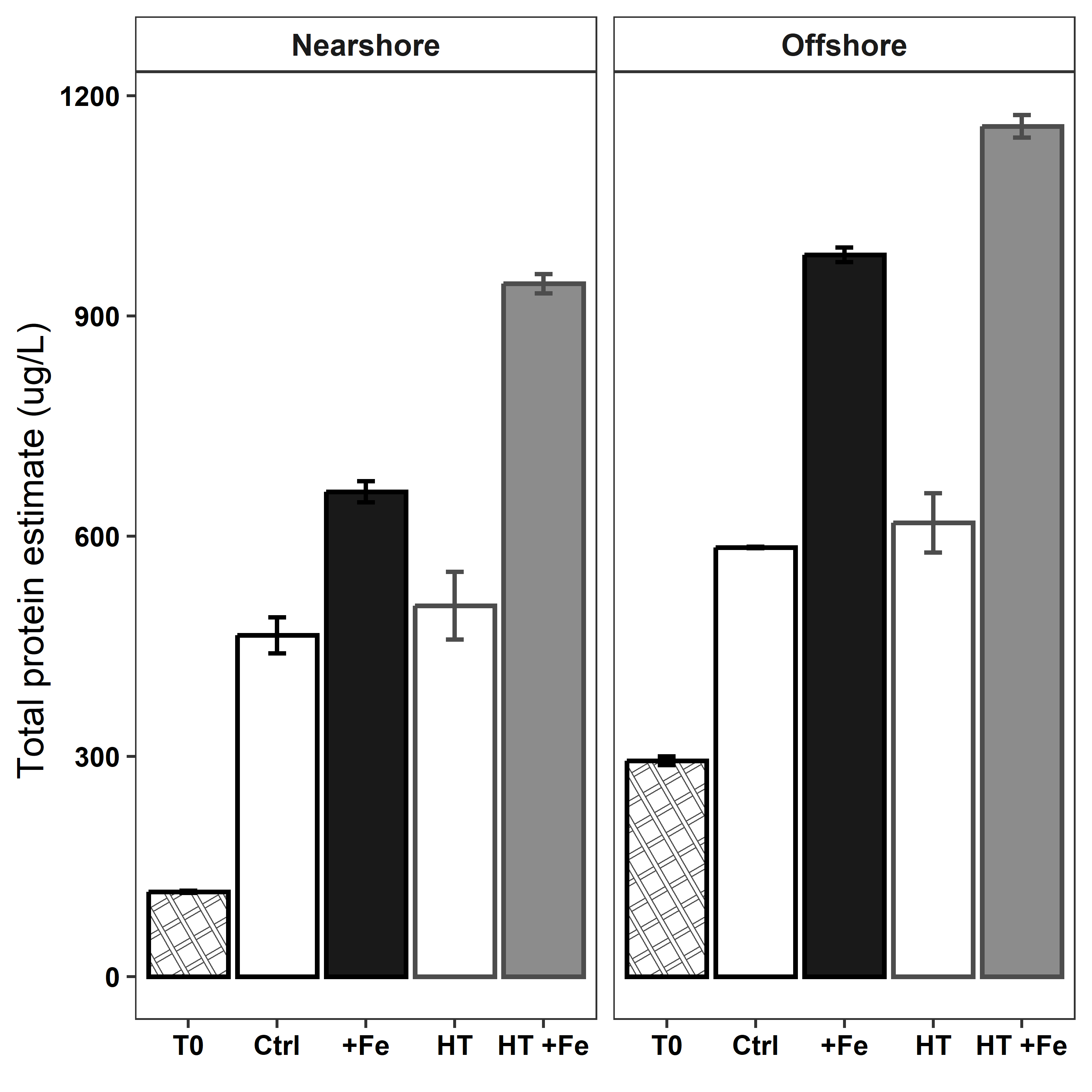

**Figure S4 –** Total protein concentrations calculated from particulate nitrogen measurements using PON*4.78 following^2^. Iron addition increases total protein synthesis, concurrent iron addition and warming cause a synergistic increase in total protein synthesis.

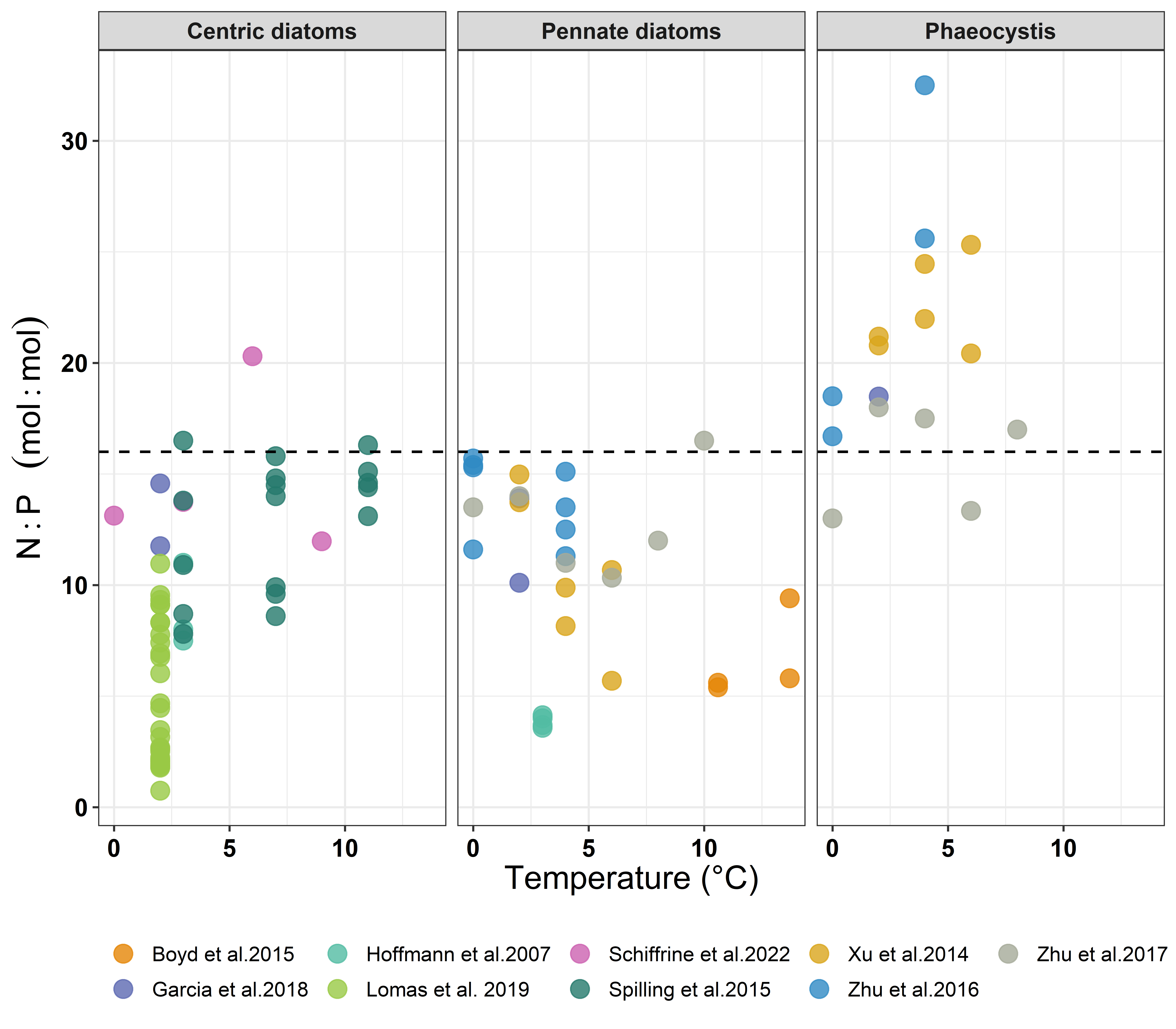

**Figure S5 –** A color coded version of Figure dD, showing N:P ratios acquired from previously published studies of laboratory cultures grown under nutrient replete conditions and harvested during the exponential phase in semi-continuous or batch growth. Centric diatoms include *Thalassiosira* sp., *Chaetoceros* sp., *Skeletonema* sp., and *Odontella* sp. Pennate diatoms include *Pseudo-nitzschia* sp. and *Fragilariopsis* sp. Data were retrieved from the cited studies either directly from text and tables, or from figures using the web tool WebPlotDigitizer^3^. Horizontal dashed horizonal lines represent Redfield N:P ratio of 16. The studies used were as follows: Boyd et al. 2015^4^, Garcia et al. 2018^5^, Hoffmann et al. 2007^6^, Lomas et al. 2019^7^, Schiffrine et al. 2022^8^, Spilling et al. 2015^9^, Xu et al. 2014^10^, and Zhu et al. 2016, 2017^11,12^.

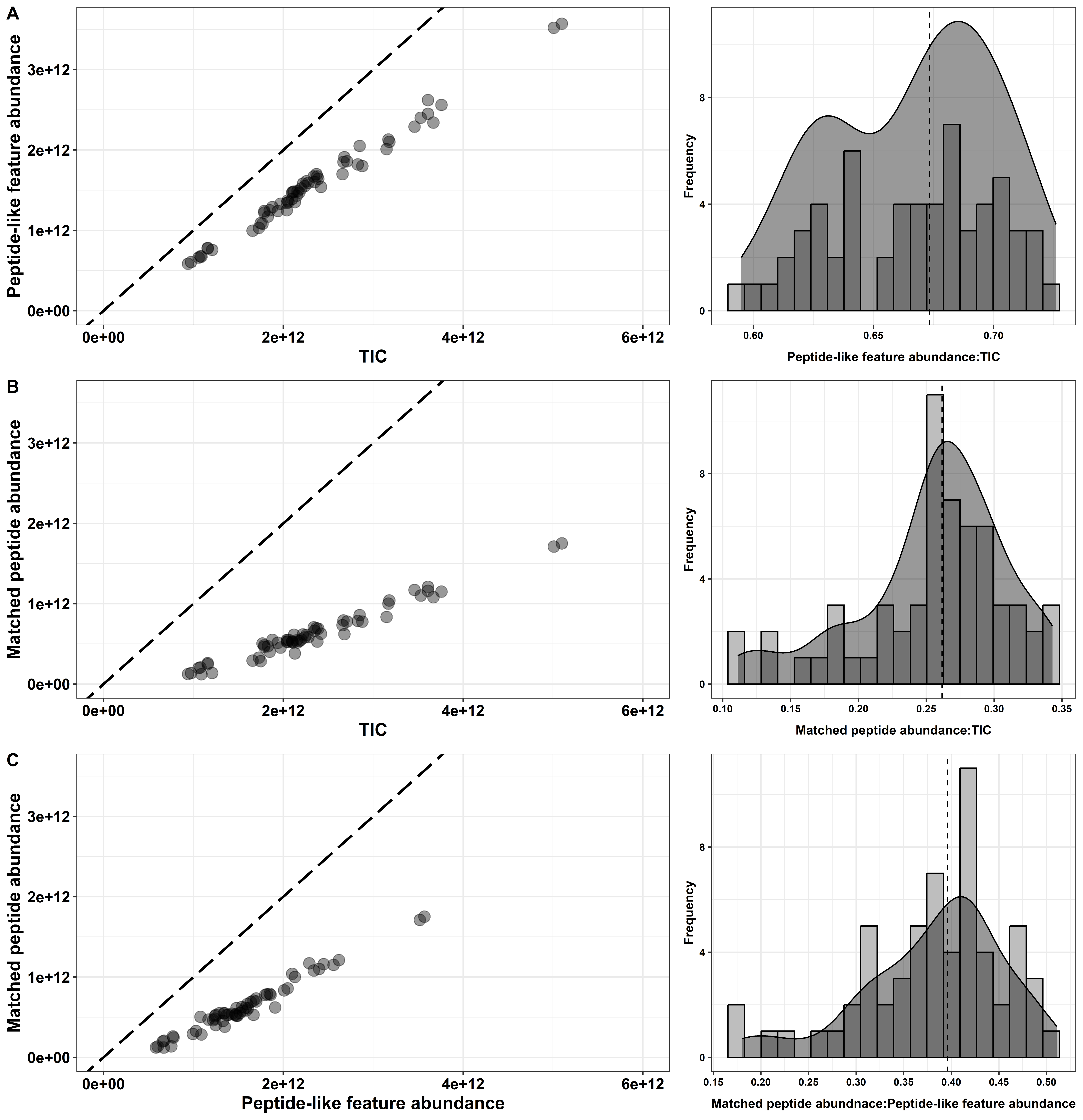

**Figure S6 – A)** Peptide-like feature abundance plotted against TIC. Frequency histogram shows the number of injections (y-axis) with a given peptide-like feature abundance: TIC ratio (x-axis), calculated up to 4 significant digits. **B)** Matched peptide abundance (database dependent) plotted against TIC. Frequency histogram shows the number of injections with a given Matched peptide abundance: TIC ratio, calculated up to 4 significant digits. **C)** Matched peptide abundance plotted against peptide-like feature abundance. Frequency histogram shows the number of injections with a given matched peptide abundance: peptide-like feature abundance ratio, calculated up to 4 significant digits. In A, B, and C, each point represents one sample injection. Diagonal dashed lines indicate 1:1 line. Vertical dashed lines indicate the median ratio.

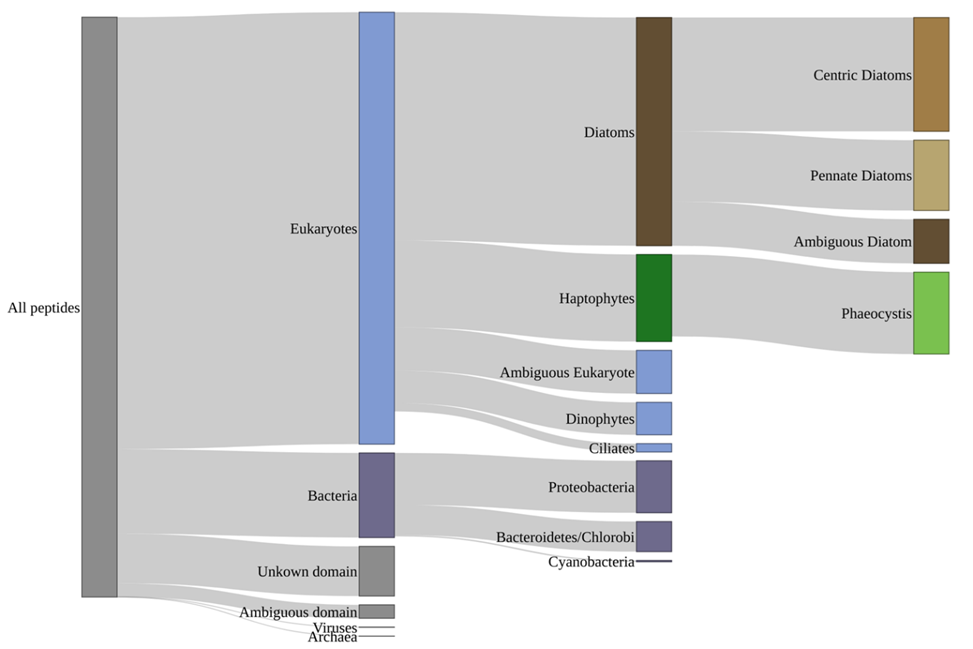

**Figure S7** – From left to right, bar height represents the number of unique peptides assigned to each taxonomic resolution. Note that the number of unique peptides here does not reflect peptide abundance. The number of unique peptides assigned to the main taxonomic groups are as follows: All peptides = 37472, Eukaryotes = 27910, Bacteria = 5466, Unknown domain (peptides that could not be linked to a taxonomic affiliation) = 3208, Diatoms = 14746, Haptophytes = 5622, Centric diatoms = 7353, Pennate Diatoms = 4547, Ambiguous diatoms (diatom peptides that could not be differentiated between pennate and centric origins) = 2846, Phaeocystis = 5287.

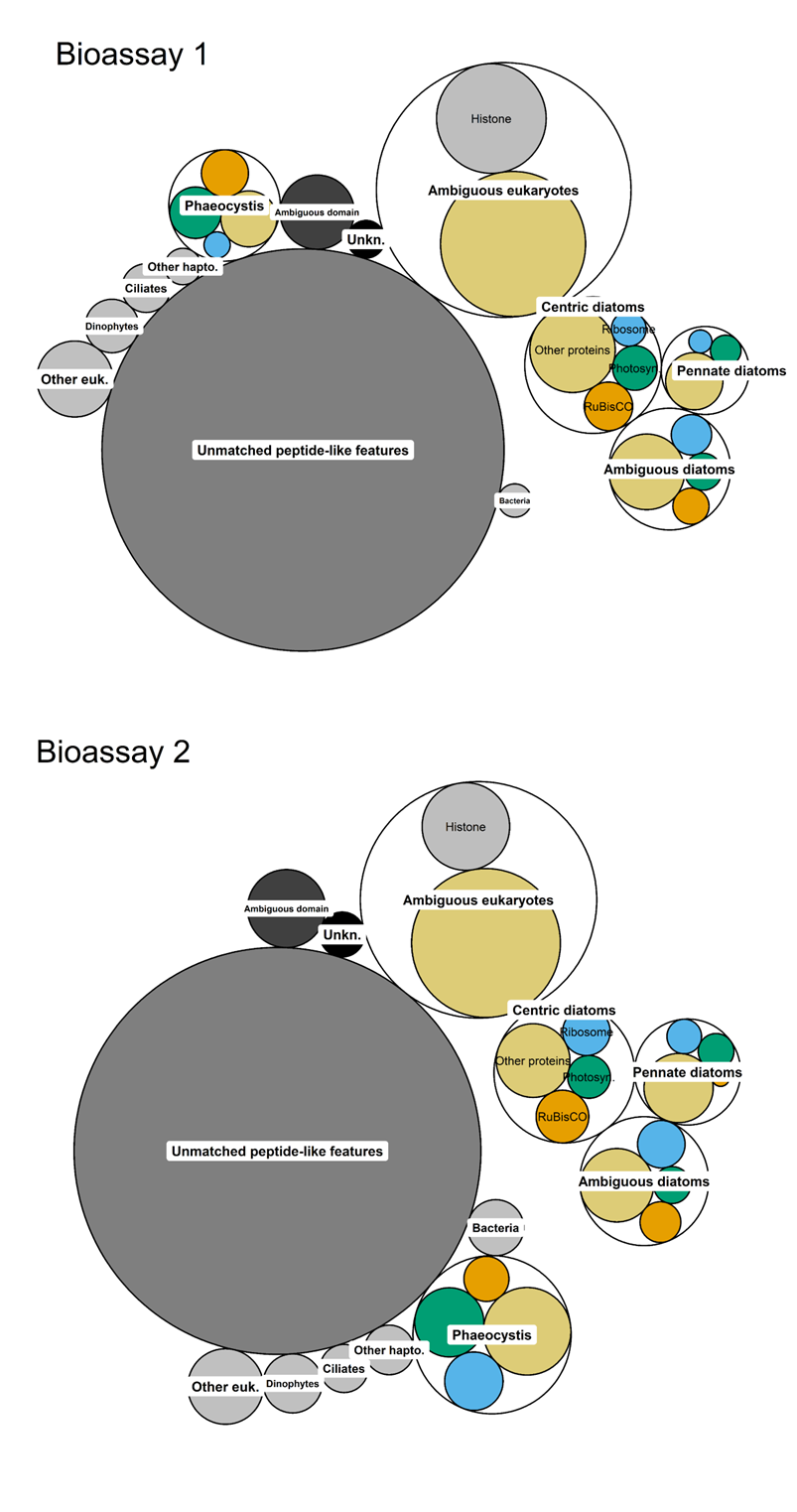

**Figure S8 –** Average abundance of major taxa and functional protein groups at T0 in Bioassay 1 and Bioassay 2. Circle size represents taxon or functional protein abundance normalized to total protein in each bioassay. “Unmatched peptide-like features” indicates the abundance of possible peptide-like features that were not matched using our database. “Unkn.” indicates matched proteins with no known function or taxonomic affiliation.

**Supplemental Tables**

**Table S1 *–*** Liquid chromatography flow rates and solvent composition settings. Arrows indicate linear increase or decreases in Solvent B* and A** composition. p

| **Time (minutes)** | **Flow (μL/min)** | **% Solvent B** | **% Solvent A** |
| --- | --- | --- | --- |
| 0 - 15 | 0.3 | 5 | 95 |
| 15.1 - 90 | 0.25 | 5 🡪 30 | 95 🡪 70 |
| 90.1 - 102 | 0.25 | 30 🡪 55 | 70 🡪 45 |
| 102.1 - 106 | 0.3 | 55 🡪 95 | 45 🡪 5 |
| 106.1 - 110 | 0.3 | 95 | 5 |
| 111 - 125 | 0.3 | 5 | 95 |

* Solvent A: H_2_O, 0.1% Formic Acid

** Solvent B: Acetonitrile, 0.1% Formic Acid

**Table S2 –** Q-Exactive hybrid quadrupole-Orbitrap mass spectrometer settings.

| **Parameter** | **Setting** |
| --- | --- |
| TopN | 8 |
| Intensity Threshold | 8.3e4 |
| Dynamic Exclusion | 30 seconds |
| MS1 Scan Resolution | 140000 |
| MS1 Scan Range | 400 to 2000 m/z |
| MS1 Automatic Gain Control Target | 3e6 |
| MS2 Scan Resolution | 17500 |
| MS2 Scan Range | 200 to 2000 m/z |
| M2 Automatic Gain Control Target | 1e6 |
| MS2 Isolation Window | 2.0 m/z |
